# An explainable graph neural network approach for effectively integrating multi-omics with prior knowledge to identify biomarkers from interacting biological domains

**DOI:** 10.1101/2024.08.23.609465

**Authors:** Rohit K. Tripathy, Zachary Frohock, Hong Wang, Gregory A. Cary, Stephen Keegan, Gregory W. Carter, Yi Li

## Abstract

The rapid growth of multi-omics datasets, in addition to the wealth of existing biological prior knowledge, necessitates the development of effective methods for their integration. Such methods are essential for building predictive models and identifying disease-related molecular markers. We propose a framework for supervised integration of multi-omics data with biological priors represented as knowledge graphs. Our framework leverages graph neural networks (GNNs) to model the relationships among features from high-dimensional ‘omics data and set transformers to integrate low-dimensional representations of ‘omics features. Furthermore, our framework incorporates explainability methods to elucidate important biomarkers and extract interaction relationships between biological quantities of interest. We demonstrate the effectiveness of our approach by applying it to Alzheimer’s disease (AD) multi-omics data from the ROSMAP cohort, showing that the integration of transcriptomics and proteomics data with AD biological domain network priors improves the prediction accuracy of AD status and highlights functional AD biomarkers.

## Introduction

Advances in high-throughput technologies have led to an explosion in the generation and availability of molecular data, encompassing the analysis of diverse biomolecules such as DNA, RNA, proteins and metabolites [1]. This has, consequently, enabled the study of fundamental processes such as gene expression [2] and DNA methylation [3], and opened new avenues for understanding complex biological systems and disease mechanisms. Profiling multiple ‘omics modalities in a disease cohort can provide a more comprehensive understanding of how distinct molecular processes operate in tandem to contribute to disease development and progression. Deriving such insights necessitates the development of methods for multimodal integration. Indeed, suitably designed integrative analysis can not only improve predictive outcomes but also help identify novel therapeutic targets, enabling the development of personalized medicine [4].

Integrating and analyzing multi-omics datasets poses significant computational challenges. These datasets are typically high-dimensional and heterogeneous, making data reduction and identification of shared patterns essential. Additionally, omics coverage may be incomplete, leading to missing data and potential biases. A variety of unsupervised methods have been proposed to address these challenges and derive insight from multi-omics datasets. The standard approach to dealing with the high-dimensionality of multi-omics data is to employ matrix factorization techniques. Methods such as MOFA (multi-omics factor analysis) [5], iCluster [6] and iNMF [7] look for latent factors shared across data modalities. Another class of unsupervised methods attempts to produce a unified representation of heterogeneous ‘omics modalities by clustering samples based on similarities shared between their omics profiles – see, for instance, similarity network fusion (SNF) [8] and unsupervised graph kernel learning approaches [9, 10].

In spite of their applications to a variety of bulk and single-cell multi-omics datasets for discovering molecular mechanisms and identifying biomarkers [11], unsupervised methods do not allow one to detect signals or patterns pertinent to a specific target phenotype, such as a particular disease of interest. Meanwhile, methods for integrating heterogeneous data in the supervised setting are relatively sparse, where the challenge posed by the high-dimensionality of multi-omics data is further compounded by the small dataset size (i.e., the number of patient samples is significantly smaller than the total number of biological molecules profiled); particularly in the bulk ‘omics setting. Existing methods for supervised integration seek to exploit structures in ‘omics datasets: patients with similar ‘omics profiles are likely to share similar disease diagnoses. Based on this principle, several methods have been proposed to leverage graph neural networks (GNNs) to pose the task of patient phenotype prediction as a graph node classification problem. MOGONET [12] leverages GNN feature extractors by using empirically generated patient similarity networks. MoGCN [13], on the other hand, learns a unified GNN model using a patient similarity graph topology generated with SNF and low-dimensional node features learnt through an autoencoder. While methods based on patient-similarity structures can alleviate the computational challenges associated with high-dimensionality in data features and low sample size, they do not leave any room for exploiting structures in the feature space, i.e. prior information about the relationship between biomolecules being measured.

In this work, we propose a novel explainable GNN framework, or GNNRAI (<u>GNN</u>-derived <u>r</u>epresentation <u>a</u>lignment and <u>i</u>ntegration), for supervised integration of multi-omics data. Unlike existing methods, such as MOGONET and MoGCN, which use networks to model relationships among samples, we use graphs to model relationships among modality features (for example, genes in transcriptomics and proteins in proteomics data). This enables us to encode prior biological knowledge as graph topology. Given *k* ‘omics modalities, each sample is represented as *k* graphs in our framework. We leverage supervised GNNs to learn modality-specific low-dimensional graph embeddings. These low-dimensional embeddings are first aligned to each other to enforce shared patterns, and then integrated using a set transformer [14]. The integrated multi-omics representations are used to predict the target phenotype. Our model architecture allows us to incorporate samples with incomplete ‘omics measurements and avoid a reduction in statistical power. To identify predictive modality features, we employ the method of integrated gradients [15] which estimates the importance of each feature to model predictions. We demonstrate the effectiveness of our framework by applying it to the task of predicting Alzheimer’s disease (AD) status by integrating transcriptomics and proteomics data from the Religious Order Study/Memory Aging project (ROSMAP) cohort. Given that proteomics data typically have a much smaller number of features relative to transcriptomics data, exacerbated by the smaller number of samples with proteomic data in the ROSMAP cohort, multi-omics integration methods might mask the role of the proteomics modality [16]. Our results show that proteomics data are more predictive than transcriptome data in the ROSMAP cohort and the integration of the two data modalities using our GNNRAI improves upon the predictive performance of the two unimodal models. Graph topology for our ‘omics-specific GNNs is derived from recent work on AD biological domains (biodomains, or BDs), which are expertly curated knowledge graphs for AD-associated endophenotypes [17]. Our modeling framework is compared to the MOGONET approach on held-out validation data and shows improved prediction metrics. We derive important ‘omics features within AD-associated biodomains via the integrated gradients method. Due to the incorporation of prior biological pathway knowledge in our GNNRAI, the identified biomarkers are predominantly functional. Finally, we probe our trained single-biodomain models to derive interactions between these biodomains using a set transformer in a second modeling stage. Interpretation of biodomain interactions via the method of integrated Hessians [18] allows us to map how AD biodomains potentially interact to drive disease in combination.

## Results

### GNNRAI for supervised multi-omics integration and biomarker identification

In this work we developed an AI framework, GNNRAI (GNN-derived representation alignment and integration), for performing supervised multi-omics data integration, accommodating potentially incomplete data and identifying informative biomarkers and biological interactions. The backbone of our proposed method consisted of GNN-based feature extractor modules. Omics data, coupled with prior knowledge graphs, were processed through these GNN-based feature extractors to produce low-dimensional embeddings. Modeling the relationships between markers reduced the training sample size burden since correlation structure reduces the effective dimensions in high-dimensional omics data. Leveraging prior pathway knowledge and integrating multi-omics data maximized the likelihood that the identified informative features were functional.

A schematic of our end-to-end GNNRAI model is shown in Figure 1. The MLP classifiers were designed for samples with a single modality, while the set transformer module was used exclusively for samples with complete multi-omics measurements. This architecture facilitated efficient training on incomplete multi-omics datasets, as the feature extractor modules were updated by all samples regardless of the completeness of their omics data. To explain the predictions from our model, we leveraged the integrated gradients method [15], a method for *post-hoc* interpretability of black-box models. Furthermore, we used the method of integrated Hessians [18] to extract informative biological interactions between single-biodomain model representations. Though we only demonstrated the integration of two modalities, it is straightforward to extend to multiple modalities.

**Figure 1:**
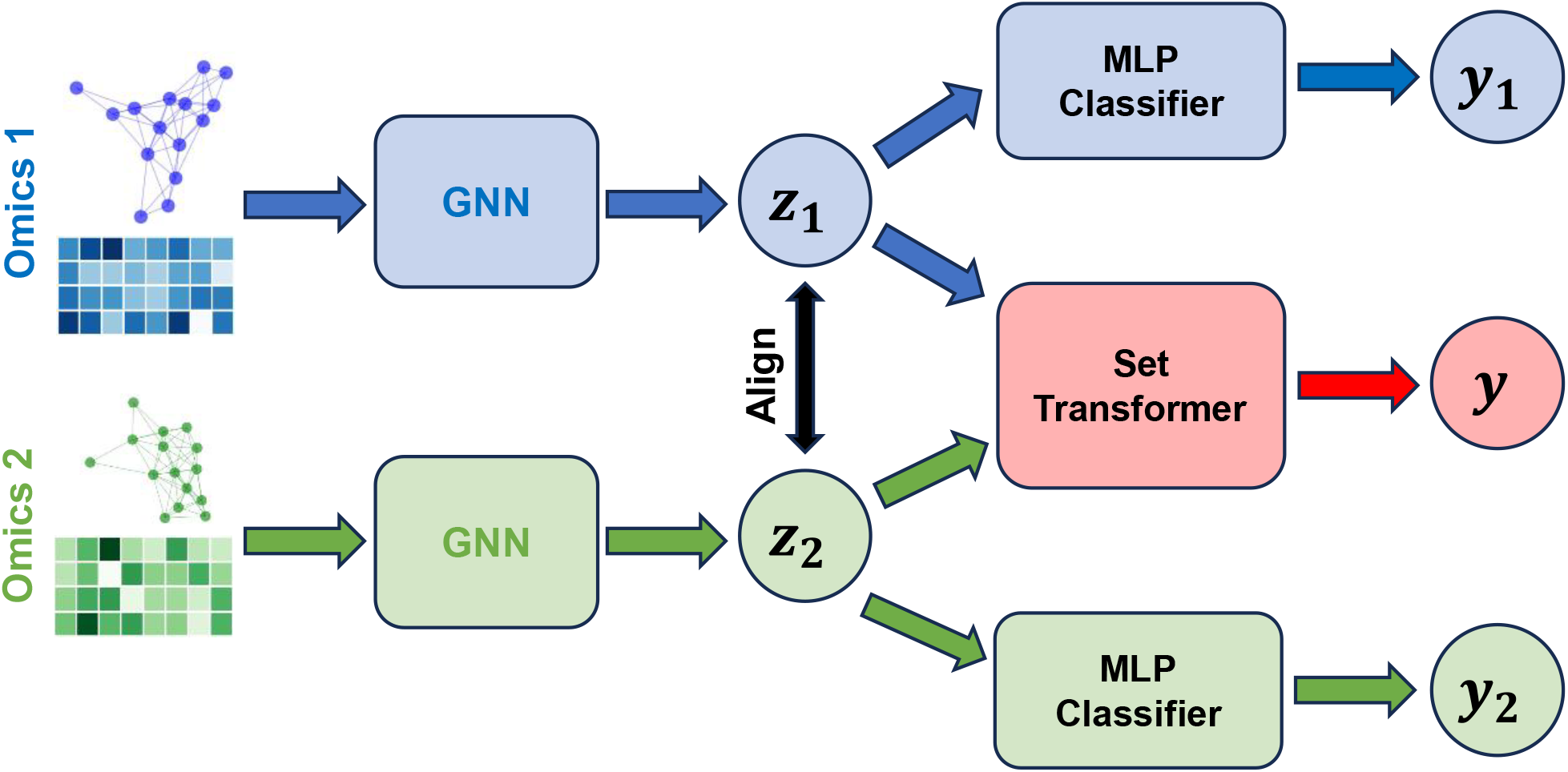
Schematic of our end-to-end integrative model GNNRAI. Data from individual ‘omics modalities are processed in their respective GNN feature extractors to produce low-dimensional embeddings (z_1_ and z_2_). z_1_ and z_2_ are then aligned and integrated through a set transformer. They are also processed through separate MLP (multi-layer perceptron) classifiers to produce modality-specific predictions of the target when a sample has incomplete multi-omics measurements.

### Alzheimer’s disease patient classification datasets

In our study, we implemented the GNNRAI framework to integrate transcriptomic and proteomic data for the binary classification of Alzheimer’s disease within the ROSMAP cohort. We analyzed gene and protein data from the dorsolateral prefrontal cortex (DLPFC) brain region. The processing of the data and the criteria for AD diagnosis were elaborated in the Methods section. Building on Cary’s 2024 research on AD biodomains [17], we created 16 datasets for AD classification, each representing a different biodomain. These datasets were complemented with knowledge graphs derived from querying the Pathway Commons database. Refer to the Methods section for details on biodomains and graph sizes for each biodomain. After data processing, we had 228 samples with both transcriptomic and proteomic data, 59 with only proteomic data, and 336 with only transcriptomic data. Our GNNRAI models were trained on each of these 16 biodomain-specific datasets. The datasets consisted of graphs with nodes representing genes or proteins from a biodomain, with their expression or abundance values as node features, structured by the biodomain’s knowledge graph from querying the Pathway Commons [19]. Each sample was labeled with a binary indicator to denote whether it was from an AD patient or a healthy control.

### Proposed GNNRAI AI framework outperformed benchmark MOGONET method on AD/control classification

The *multi-omics graph convolutional network*, or MOGONET [12], was a supervised learning framework for integrating multi-omics data using GNNs. MOGONET processed individual modalities separately by constructing patient similarity networks using the cosine distance metric to assign edges. Graph neural networks operated on these patient similarity networks to make modality-specific predictions, which were then integrated through a view correlation discovery network (VCDN). In contrast to MOGONET, our approach imposed a network topology over the space of input features within each modality. The MOGONET architecture made it implausible to incorporate priors on the space of features (such as AD biodomains). Furthermore, MOGONET required samples to have complete measurements (i.e., no missing modalities). We trained our unimodal and integrative models on the set of samples with both transcriptomics and proteomics measurements for each of the 16 BDs and compared their validation predictive performance to that of MOGONET trained on the same datasets. A comparison of the validation performance between these models is shown in Figure 2. We observed that when trained on an equal number of samples, unimodal proteomics models consistently outperformed unimodal transcriptomics models. Despite having fewer features (see Table 3), proteomics data were more predictive in the ROSMAP cohort, aligning with [20]. Our integrative models outperformed the integrative MOGONET models in 13 out of 16 BD datasets (except for *apoptosis, Tau homeostasis* and *vasculature)*. Additionally, for seven BD datasets (*cell cycle, endolysosome, immune response, lipid metabolism, metal binding, oxidative stress* and *proteostasis*), our unimodal proteomics models surpassed the multimodal MOGONET models. This was likely because proteomics and transcriptomics data were not always consistent, and MOGONET integrated modality-specific predictions rather than modality representations. In contrast, our framework’s integration of transcriptomics and proteomics modalities improved the unimodal predictive performance across all 16 BD datasets, demonstrating effective integration.

**Figure 2:**
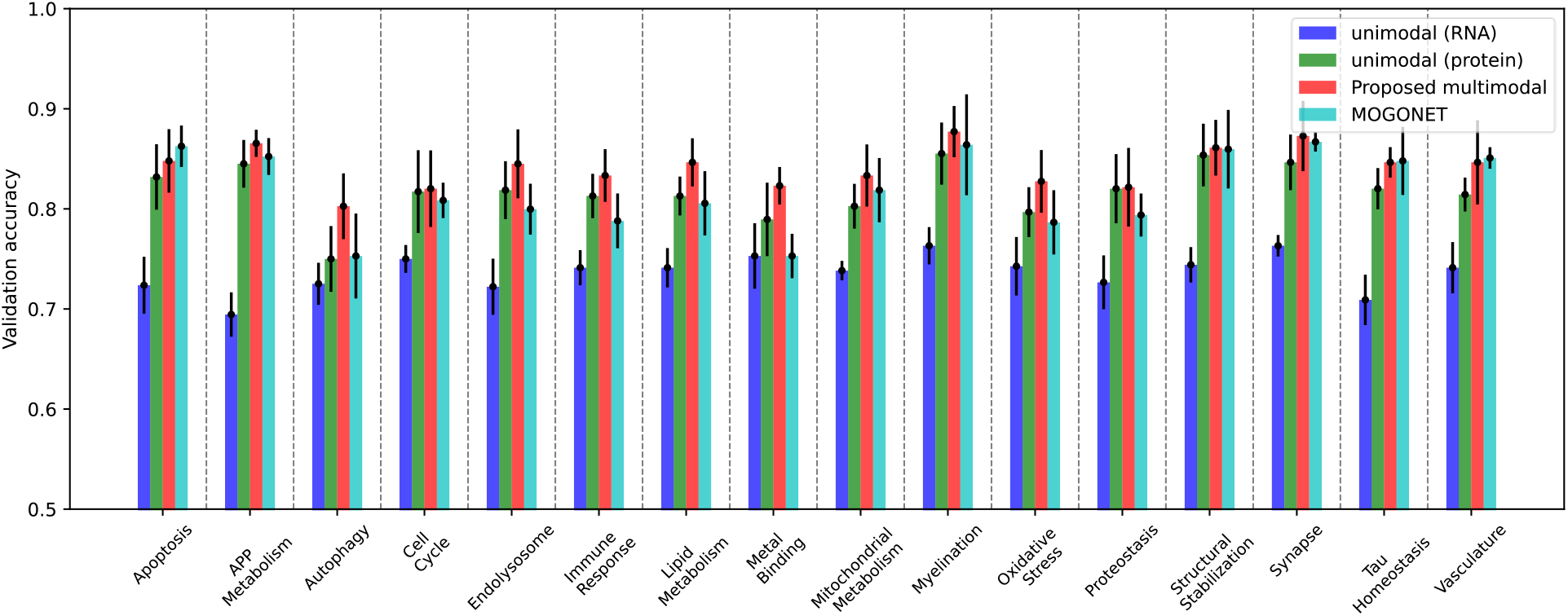
Validation performance of our proposed integrative model (red) compared to validation performance of the benchmark MOGONET model (cyan) on the set of common saples having both proteomics and transcriptomics measurements. The performance of the integrative models is also compared to unimodal GNN transcriptomics (blue) and proteomics (green) models.

### Multi-omics GNNRAI models outperformed unimodal GNNRAI models trained on transcriptomics and proteomics alone

For samples with complete measurements, our model’s performance was evaluated using the held-out validation set predictions from the integrative component (i.e., the set transformer in Figure 1). For samples with incomplete measurements, predictions were based on their respective unimodal classifiers. We found that integrating the two modalities resulted in better performing classifiers compared to the unimodal counterparts. This finding was significant given that we had 564 samples with transcriptomic data but only 287 with proteomic data. A larger number of less predictive transcriptomic samples could obscure the superior performance of proteomic samples if the useful information from both modalities was not effectively aligned and integrated.

To ensure a fair comparison, we used two sets of validation samples for evaluating the performance. The first validation dataset comprised samples with transcriptomics measurements (used to test the unimodal transcriptomics GNN models). The second validation dataset included samples with proteomics measurements (used to test the unimodal proteomics models). Figure 3 compares the validation accuracy of GNN models trained solely on transcriptomics and proteomics samples with the end-to-end multimodal models trained on all samples. Validation performance for unimodal models were denoted as ‘unimodal-RNA’ and ‘unimodal-prot’, while the two validation scores from the integrative model were denoted as ‘integrated-RNA’ and ‘integrated-prot’ respectively. In spite of the smaller training dataset, ‘unimodal-prot’ consistently outperformed ‘unimodal-RNA’ across all 16 BDs, reiterating that proteomics data provided more AD-predictive information than transcriptome data in the ROSMAP cohort. When we compared the multimodal to unimodal performance, ‘integrated-RNA’ consistently surpassed ‘unimodal-RNA’ across all 16 BDs, demonstrating that integrating proteomics with transcriptomics data enhanced classification performance. Similarly, there was a consistent performance improvement from ‘unimodal-prot’ to ‘integrated-prot’, albeit less pronounced than in the RNA modality. Furthermore, ‘integrated-prot’ was generally better than ‘integrated-RNA’ except for *lipid metabolism* BD, despite the fact that the proteome-specific classifier was trained on only 59 samples compared to 336 samples for the transcriptome-specific classifier. This suggests that the target-predictive signals from transcriptomic and proteomic embeddings were aligned and integrated effectively, resulting in smaller performance differences between samples with transcriptomic data and proteomic data than in unimodal transcriptomic models.

**Figure 3:**
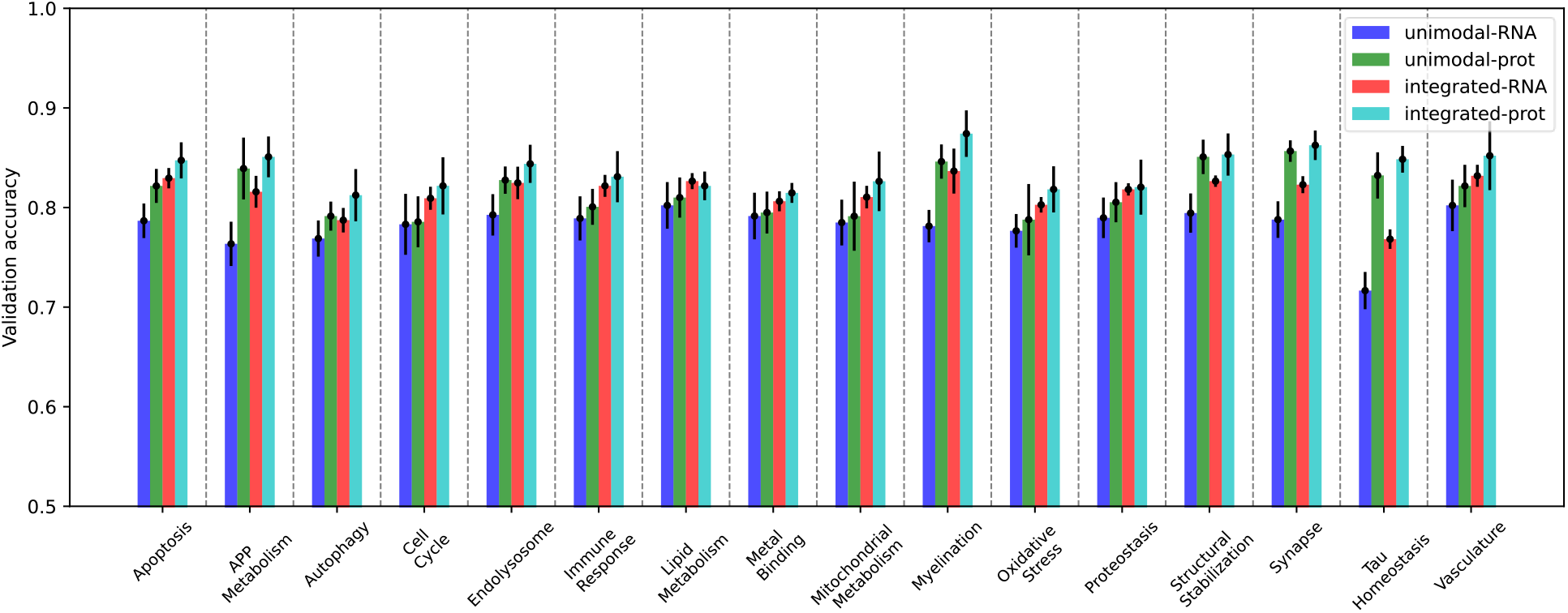
Validation performance of the integrative multimodal model (red and cyan) compared to performance of unimodal GNN models (blue and green) trained on the incomplete multi-omics datasets for 16 AD biodomains. The performance of the multimodal model is calculated on two sets of validation samples – the set of all validation samples with transcriptomics measurements and that with proteomics measurements.

### Validation on ROSMAP, Mount Sinai Brain Bank (MSBB) and Mayo Clinic transcriptomics and proteomics data

To validate the predictive ability of our models trained on ROSMAP DLPFC samples, we curated samples with transcriptomics and/or proteomics measurements from the following studies and brain regions -

1. ROSMAP samples from the *anterior cingulate cortex* (ACC) and *posterior cingulate cortex* (PCC) brain regions with transcriptomic measurements.
2. MSBB [21] samples from the *parahippocampal gyrus* (PHG), *frontal pole* (FP), *inferior frontal gyrus* (IFG), and *superior temporal gyrus* (STG) brain regions. Only PHG tissue had both transcriptomic and proteomic data, while the remaining tissues had transcriptomics only.
3. Mayo Clinic [22] samples from the *temporal cortex* (TCX) brain region. Although TCX tissue had both transcriptomic and proteomic data, the proteomics measurements were acquired by label-free quantification, different from the tandem mass tag (TMT) quantification platform used in ROSMAP and MSBB. Hence, we did not validate ROSMAP-derived models on Mayo proteomics data.

Table 1 shows the sample counts of the curated validation datasets. The procedure for annotating MSBB and Mayo samples with ground truth labels were described in the Methods section. We noted that MSBB and ROSMAP adopted similar diagnostic criteria, while Mayo clinic cohort followed Mayo neurologist guidelines [23].

**Table 1:** Transcriptomics and proteomics sample counts in validation data.

| tissue | ROSMAP<br>(ACC) | ROSMAP<br>(PCC) | MSBB<br>(PHG) | MSBB<br>(FP) | MSBB<br>(IFG) | MSBB<br>(STG) | Mayo<br>(TCX) |
| --- | --- | --- | --- | --- | --- | --- | --- |
| RNA | 354 | 332 | 122 | 106 | 106 | 108 | 62 |
| protein | - | - | 122 | - | - | - | - |

We first applied the ROSMAP DLPFC-trained transcriptomics model to transcriptomics data from ROSMAP ACC and PCC brain regions, MSBB PHG, FP, IFG and STG brain regions and Mayo TCX brain region. For comparison, we also included the predictive accuracy on the ROSMAP DLPFC validation dataset (Figure 4). Figure 4 shows that the same GNN model has different predictive performance on different brain regions. Generally, ROSMAP PCC had slightly higher predictive accuracy than DLPFC, which was in turn higher than the predictive accuracy of ACC. For MSBB, PHG had the highest predictive accuracy, followed by IFG and STG, which predicted better than FP. MSBB PHG had the highest predictive accuracy across the three cohorts. MSBB IFG and STG had comparable performance with ROSMAP PCC, while MSBB FP had comparable performance with ROSMAP DLPFC, which predicted better than Mayo TCX. ROSMAP ACC had the lowest predictive accuracy on average. Nevertheless, ROSMAP ACC predictive accuracy was above 0.6 across the 16 AD biodomains, better than a random guess. The different predictive performance on transcriptomic data from different brain regions might be explained in terms of neuropathological burdens – the FP and DLPFC regions are impacted at a similar disease stage, whereas the PHG tends to be affected much earlier in disease progression. This could be further exacerbated by the differences in cohort sample selection between ROSMAP and MSBB. MSBB samples were selected for multi-omics profiling based on the presence of remarkable AD neuropathology, whereas the ROSMAP study was a longitudinal cohort and did not pre-select samples for extreme neuropathology. Thus, disease signatures learned in the ROSMAP DLPFC samples could be amplified in the MSBB samples, leading to an increase in the predictive accuracy of our models. This also implied that transcriptomic signatures might be translational across relevant brain tissues.

**Figure 4:**
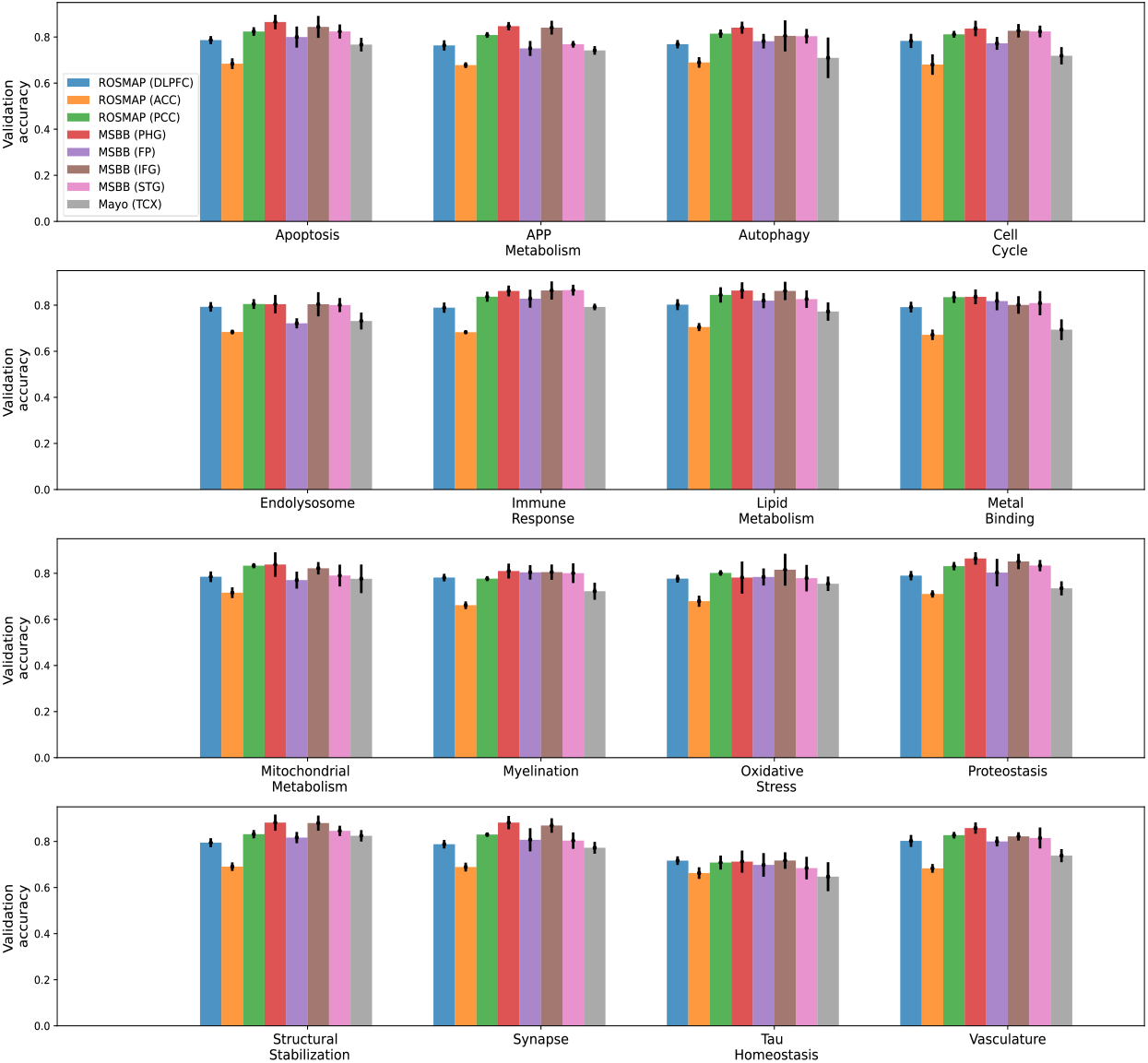
Predictive performance of applying unimodal transcriptomics model trained on ROSMAP DLPFC training dataset to validation transcriptomics samples from various cohorts and brain tissues.

Next, we applied the ROSMAP DLPFC-trained proteomics and integrative models to proteomics and multi-omics data from MSBB PHG brain region respectively (Figure 5). We also included ROSMAP DLPFC and MSBB PHG transcriptomics performance in Figure 5 for comparison. MSBB PHG had lower predictive accuracy than ROSMAP DLPFC for proteomics and integrative models. Unlike in ROSMAP, where proteomics data had higher predictive accuracy than transcriptomics data and the integrative model improved upon the two unimodal models, proteomics data were less predictive than transcriptomics data, and the integrative model had lower predictive accuracy than the proteomics model for 9 AD biodomains in MSBB. This might be due to the fact that the GNN models were trained on a much smaller set of ROSMAP proteomics data (287×2/3=191 samples) than ROSMAP transcriptomics data (564×2/3=376 samples), causing poor generalization performance on unseen data.

**Figure 5:**
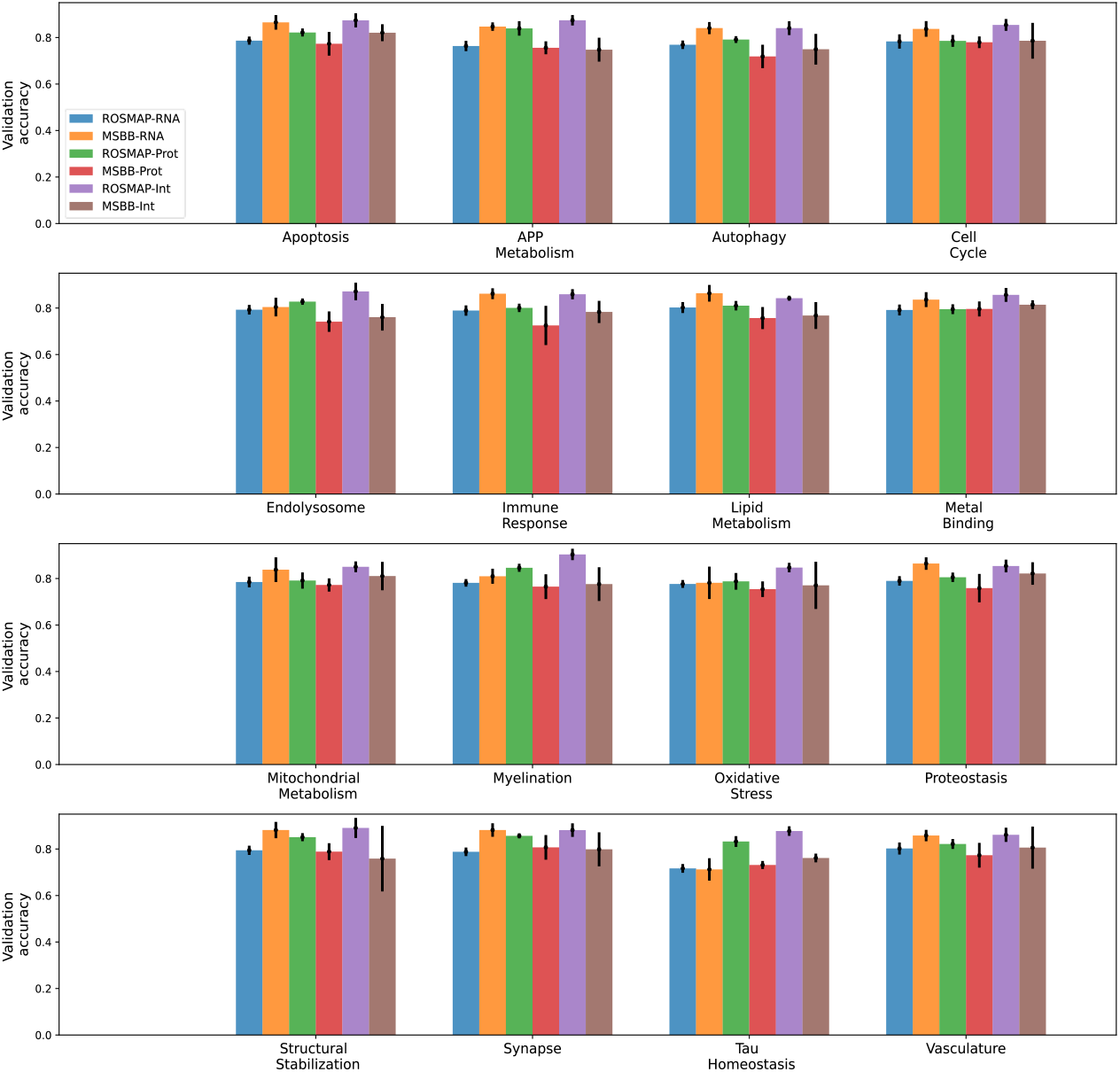
Predictive performance of applying unimodal and multimodal models trained on ROSMAP DLPFC training dataset to samples from ROSMAP DLPFC validation and MSBB PHG validation datasets. Prot: protein; Int: integrative.

### Identification of biomarkers relevant to AD

We applied the integrated gradients method to our trained multi-omics GNNRAI models to derive importance scores on input graph nodes (genes and proteins). We used a permutation-based approach to determine the importance score threshold by controlling the false discovery rate (FDR) to be below 0.05. Like standard permutation procedure for multiple hypothesis testing, we treated the original importance scores as the observed test statistics, generated 300 permuted datasets by randomly permuting the training labels and trained our integrative models on the permuted datasets. The resulting models were called null models. The importance scores for all genes/proteins in a graph from each sample of the 300 null models were used as background test statistics. Since we calculated FDR rather than corrected *p*-values, the estimated empirical FDR was confidently accurate for B=100-200 [24]. See Methods for the procedure to calculate permutation-based FDR.

We determined whether a gene/protein was informative only in correctly predicted validation samples. The top 20 AD-predictive genes/proteins are shown in Table 2. To rank the informative genes/proteins identified across the analyses, we added the number of unique sample identifiers for which a given gene/protein was identified as informative across modalities and 16 AD biodomains for each study (columns of Total samples in Table 2), divided by the number of correctly predicted validation samples in each study to obtain the fraction of samples for each gene/protein where it is informative, then calculated the average fraction across studies, based on which genes/proteins were ranked.

**Table 2:** Top 20 informative genes across studies.

| rank | gene/<br>protein | avg fraction<br>of samples | total samples<br>(ROSMAP DLPFC) | total samples<br>(MSBB PHG) |
| --- | --- | --- | --- | --- |
| 1 | MDK | 0.84 | 118 | 65 |
| 2 | APP | 0.672 | 109 | 43 |
| 3 | NTN1 | 0.62 | 92 | 45 |
| 4 | GFAP | 0.569 | 67 | 52 |
| 5 | ICAM1 | 0.541 | 66 | 48 |
| 6 | LTF | 0.532 | 67 | 46 |
| 7 | FLNA | 0.466 | 33 | 56 |
| 8 | CD44 | 0.46 | 43 | 49 |
| 9 | APOE | 0.459 | 54 | 42 |
| 10 | IL1B | 0.42 | 76 | 22 |
| 11 | VGF | 0.415 | 52 | 36 |
| 12 | FLT1 | 0.39 | 68 | 22 |
| 13 | PTN | 0.382 | 48 | 33 |
| 14 | IQGAP3 | 0.364 | 58 | 24 |
| 15 | LGMN | 0.35 | 59 | 21 |
| 16 | GJA1 | 0.347 | 34 | 36 |
| 17 | PRKCD | 0.344 | 33 | 36 |
| 18 | GIPR | 0.34 | 58 | 20 |
| 19 | TAC1 | 0.336 | 23 | 41 |
| 20 | ADCYAP1 | 0.335 | 34 | 34 |

The top ranked gene in these analyses was *MDK*, which was informative in the binary classification task for 183 unique validation samples (average of 84% of correctly predicted validation samples). *MDK* is a secreted growth factor that has consistently been identified among a suite of matrisome proteins that associate with Aβ plaques (i.e., Module M42 in [20, 25]), and has been shown to influence the aggregation of amyloid-beta, both *in vitro* and *in vivo* [26]. Notably there are also five other proteins from the M42 module among the top 20 genes (APP, NTN1, APOE, FLT1, and PTN) and module M42 proteins are significantly enriched among the most informative genes (GSEA adjusted p-value = 0.004). *VGF* was another top-ranked gene in these analyses and has consistently been identified as a robust biomarker for AD [27, 28], as well as being a top predicted regulator of multiscale AD networks [29].

The four genes in the top 20 that had the highest integrated AD <u>T</u>arget <u>R</u>isk <u>S</u>core (TRS [17]), were *APP, APOE, LGMN*, and *LTF*. Each of these genes were informative for over 80 total samples and had TRS in the top 2% of all scored genes. *APP* is a well-known disease gene that is the proteolytic precursor of the Aβ peptide, which is a major component of one of the hallmark neuropathologies of the disease, and variants within *APP* are causal for rare autosomal dominantly inherited forms of AD. Distinct alleles of the *APOE* gene represent both the strongest genetic risk and strongest genetic protective factors in genome-wide association studies of late-onset AD [30], and other APOE alleles have been found to be suppressors of mutations causing autosomal dominant AD [31]. *APOE* has been implicated in several disease endophenotypes including the accumulation of both Aβ plaques and tau neurofibrillary tangles, mediating glial cell response, and disturbances to the blood-brain barrier that occur during AD pathogenesis [32]. *LTF*, or lactotransferrin, has recently been identified as a predictor of Aβ burden [33]. *LGMN*, also known as δ-secretase, is an asparagine endopeptidase that is involved in the cleavage of both tau [34] and *APP* [35] proteolysis, which is linked to increased pathogenicity in each case. This corresponded with the findings from the individual gene/protein analyses where *LGMN* was the most impactful in Tau Homeostasis and APP Metabolism biodomains.

The remaining eleven genes from the top 20 were novel candidate biomarkers that have not been previously linked to AD pathogenesis in published studies. However, their predictive potential for AD suggested they warrant further investigation. For example, *IQGAP3* was ranked #14 in this analysis, had a TRS in the top 10%, and was differentially expressed in both transcriptomics and proteomic samples. Despite having no publications where *IQGAP3* is implicated in AD, it was linked with cytoskeletal maintenance and neurite outgrowth [36] which is consistent with its role in the Structural Stabilization domain in these analyses.

At least nine genes among the top 20 were strongly related to AD biology. Moreover, among the six M42 module members that were ranked top 20, three (APOE, FLT1, PTN), unlike MDK, NTN1, and APP, did not show high magnitude fold changes. These demonstrated that our integrative method successfully identified functional features by incorporating prior biological pathway knowledge.

### Detecting interactions among biodomains

For a multi-modal integrative model trained on a given biodomain, the class token representation from the final set transformer was a single low-dimensional embedding that unified information across modalities within the biodomain. We, therefore, collected class token representations in integrative models from all 16 BDs, and trained an auxiliary set transformer to integrate information across biodomains. Integrated Hessians [18] was applied to this second set transformer model to derive interaction scores between its input tokens. The biodomains partition gene functions into distinct molecular endophenotypes. However, these endophenotypes can and do interact during the etiology of the disease. Therefore, a primary goal of utilizing the biodomain framework is to identify interactions between domains that could broaden our understanding of the disease development.

The top interactions detected are shown as a graph where each node is a BD in Figure 6. Interactions were ranked by repeating the model training and informative interaction identification process ten times with different random weight initializations. From each iteration, the top ten percent of interactions, determined by ranking the number of samples for which each interaction was informative, were noted. Interactions present in the top ten percent three or more times out of ten were recorded. Rank was determined first by the number of appearances in the top ten percent, then, in the case of ties, by the total number of samples in which the interaction was informative. The domain nodes with the largest degree were Lipid Metabolism (degree = 9), followed by Mitochondrial Metabolism (degree = 5), Synapse and Endolysosome (degree = 4, each). The centrality of these domains was supported by the observation that Synapse, Lipid Metabolism, and Mitochondrial Metabolism were among the top risk-enriched biodomains [17]. The observation that Lipid Metabolism was a hub in this graph suggested that aspects of Lipid Metabolism influenced many other disease processes. The centrality of Lipid Metabolism to AD pathogenesis was supported by myriad observations from recent decades, including genetic studies that implicate Lipid Metabolism associated genes (e.g. *APOE, CLU, ABCA7, SORL1*) in driving AD risk [37], the observation that amyloid-beta production occurs in lipid raft membrane microdomains [38], recent lipidomic studies that identify changes in lipid species that are specific to the disease [39, 40], and many more.

**Figure 6:**
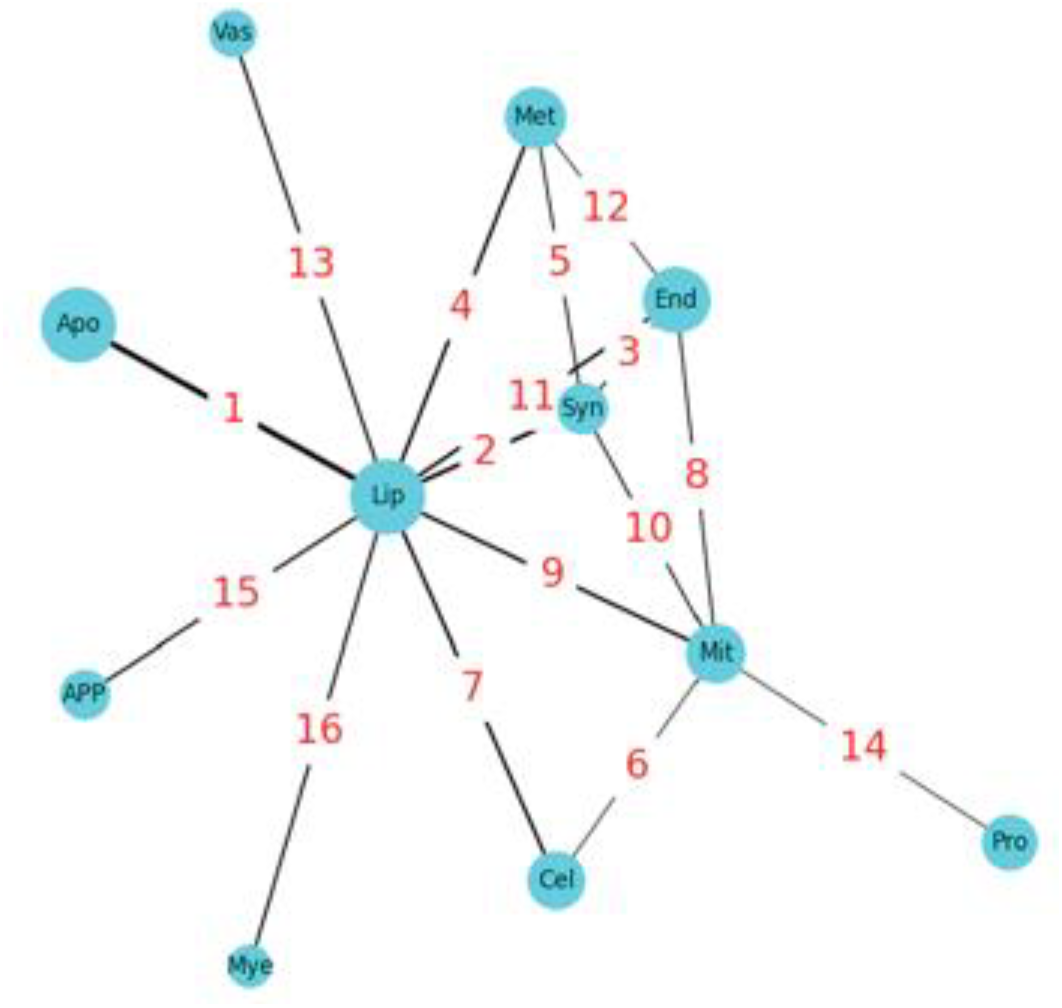
Interactions between biodomains based on Integrated Hessians analysis on multi-modal model trained on samples with complete multi-omics. The nodes represent biodomains and edges represent interactions with the edge annotations being the rank of the interaction. The biodomain names based on the 3-letter abbreviations are: apoptosis (apo), lipid metabolism (lip), synapse (syn), endolysosome (end), metal binding (met), cell cycle (cel), mitochondrial metabolism (mit), myelination (mye), vasculature (vas), proteostasis (pro) and APP metabolism (app).

Other interaction relationships represented in this graph were informative and supported by evidence from the literature. As mentioned above, the edge between APP Metabolism and Lipid Metabolism was supported by the influence of lipid rafts on amyloid-beta production. The link between Lipid Metabolism and Mitochondrial Metabolism was also very well supported given that β-oxidation of fatty acids, which is the primary catabolic pathway, occurs in the mitochondrial matrix. Given that mitochondria provide the requisite energy and precursor metabolites for cell cycle progression, and that mitochondrial biogenesis and dynamics are influenced by cell cycle regulators, the interdependence of Cell Cycle and Mitochondrial Metabolism was also well supported. The top ranked edge between Lipid Metabolism and Apoptosis evoked ferroptosis, which is an iron-dependent cell death mechanism that is distinct from apoptosis but involves the accumulation of peroxidated lipid species [41] and is the focus of newly emerging hypotheses of disease pathogenesis [42]. Further, supporting this association is the observation that 6 of the top 20 identified informative genes – i.e. *LGMN* [43, 44], *LTF* [45, 46], *CD44* [47, 48], *PRKCD* [49], *IL1B* [50, 51], and *GJA1* [52-54] – are associated with regulating aspects of ferroptosis in diverse contexts. It was noted that the immune response biodomain, which is strongly involved in AD, is absent in Figure 6. The interplay between the immune response and lipid metabolism BDs ranked seventeen in these analyses – the highest ranking interaction not included in Figure 6. The reason for this omission in our list of top pairwise interactions remains unclear and warrants further investigation which is beyond the scope of our current work.

## Discussion

In this work, we proposed an end-to-end AI framework, or GNNRAI, for supervised alignment and integration of multi-omics data with prior information expressed as knowledge graphs. Our method was based on the use of GNNs for learning low-dimensional embeddings from high-dimensional data and could accommodate samples with missing modalities. Using the ROSMAP data, we curated 16 binary classification datasets – each dataset comprising a view of the ROSMAP gene expression and protein abundance data within an AD-associated biodomain. We noted that the size of biological domains varied significantly from smallest to largest domains and therefore allowed us to test the robustness of our approach to varying input dimensionality. Our approach has validated its efficacy in the task of integrating transcriptomics and proteomics data from the ROSMAP cohort. It has outshined the benchmark MOGNET method in 13 of the 16 BDs and shown improvements over the two unimodal models for all 16 BDs. This outcome is noteworthy considering the disparity in sample sizes, with 564 transcriptomic samples and only 287 proteomic samples. The abundance of less predictive transcriptomic samples could potentially conceal the enhanced performance of the proteomic samples unless the valuable insights from both data types are properly synchronized.

The task of integrating multi-omics data is computationally challenging for several reasons. We demonstrated that the curse of dimensionality arising from the large number of ‘omics features relative to the sample size could be overcome by leveraging the correlation structure in graphs and message passing in GNNs. Additionally, we showed that the separation of feature extraction modules and set transformer-based integration allowed us to utilize samples with missing modalities – a characteristic feature of multi-omics datasets.

Our framework allows one to integrate modalities where prior information about the relationship between input features can be expressed in the form of knowledge graphs. We leveraged existing work on AD biodomains to extract network topologies for transcriptomics and proteomics modalities. Our approach did not make a distinction in the structure of the knowledge graphs used for these modalities, thereby implicitly imposing a simplified assumption that network relationships between transcripts is exactly reproduced in the proteins they code for. Furthermore, we did not incorporate data from other modalities within the ROSMAP study, such as methylomics and metabolomics, due to the current unavailability of direct prior knowledge graphs for these modalities. However, existing transcriptomic and proteomic networks can be leveraged for the construction of gene-centered methylomics and metabolomic knowledge graphs. For instance, metabolites catalyzed by the same genes/proteins may be determined to share a relationship. For genes that have an edge in a transcriptomic network, CpG sites within their regulatory regions (promotors, enhancers etc.) may be determined to share the same edge within the corresponding methylation network. Finally, we identified informative features through the model-agnostic method of integrated gradients which derived importance scores on individual graph nodes independently. Interpretation of GNN predictions can, in theory, be enhanced by using an explanation method to identify informative subgraphs, or *motifs*. While methods in this direction did exist [55], our experience was that we were unable to extract meaningful subgraph/motifs through the application of such methods. How to identify correlated informative features efficiently is one of the important future research directions for GNNs.

## Methods

### Data preprocessing

To investigate AD mechanisms, we adopted a combination of clinical and neuropathological criteria used in [20] to assign ground truth labels (AD case or control) to patients within the ROSMAP cohort. In particular, we used clinical cognitive tests, such as MMSE (the Mini-Mental State Examination [56]), or CDR (Clinical Dementia Rating) to assess dementia: MMSE score <= 24 or CDR >= 1. Neuropathological assessment of patients was conducted post-mortem using Braak staging [57] and CERAD (The Consortium to Establish a Registry for Alzheimer’s Disease) scoring [58] to reflect AD hallmarks. CERAD scores 0-3 correspond to no AD/none, possible/sparse, probable AD/moderate, and definite/frequent, respectively. The Braak score, or seven Braak stagings, classifies the severity and distribution of tau pathology in the brain. Cases with CERAD 0–1 and Braak 0–3 without dementia at last evaluation were annotated as controls (if Braak score equals 3, then CERAD must equal 0); cases with CERAD 2–3 and Braak 3–6 with dementia at last evaluation were annotated as AD.

Downloaded RNA-Seq count data were log2 transformed and corrected for age, sex and postmortem interval (PMI) covariates. Downloaded protein abundance data were log2 transformed and median zero centered per feature. Finally, age, sex and postmortem interval (PMI) were regressed out.

For validation MSBB samples, we used similar diagnostic criteria, except MMSE was replaced with CDR since MSBB did not provide MMSE information. MSBB gene expression/protein abundance data were processed similarly to ROSMAP data. In addition, patient race was regressed out.

For patients in the Mayo study, AD and controls were taken to be the reported diagnosis according to Mayo neurologist guidelines, as described in [23]. In contrast to ROSMAP and MSBB, which made use of tandem mass tag (TMT) quantification, Mayo proteomics data were acquired with label-free quantification, hence we did not validate our models on Mayo proteomics data. Mayo gene expression data were processed similarly to ROSMAP data.

### Network priors from Alzheimer’s disease biological domains

The prior biological knowledge ascribed to nodes and edges in the knowledge graphs used for the analysis was derived from publicly accessible biological databases. These graphs provided a topological organization to the biological domains (or biodomains), which were 19 AD-associated endophenotypic descriptors, such as immune response and mitochondrial metabolism [17]. The biodomains were lists of functional biological definitions describing aspects of AD, and were defined with suites of relevant Gene Ontology (GO) terms [59]. Each GO term was annotated with a set of genes, and biological processes within a domain that were enriched for composite metrics of disease risk [17] could be identified using standard enrichment procedures. We used significantly-enriched GO terms (gene set enrichment analysis adjusted p-value < 0.01 and normalized enrichment score > 1.7) – 16 of the 19 biodomains had GO terms that met these criteria – and extracted the leading edge genes from each term to seed knowledge graph generation through a pathway reconstruction pipeline. We performed a shortest path reconstruction among all risk-enriched genes for each domain using protein-protein interaction (PPI) edge annotations from the Pathway Commons database [19], version 13. Given the nonlinear relationships implicated in most biological interactions, the shortest path to connect two genes was selected for creating an edge between two nodes. For a given protein, expressed by a gene in the biodomain, an edge was derived from the larger PPI network. The final network object consisted of edges, which were the PPI, and nodes, which were the GO term-derived gene list. See Data Availability section for the website containing the network files for the 16 biodomains. A summary of the sizes of the transcriptomics and proteomics prior networks within each biodomain is shown in Table 3.

**Table 3:** Summary of AD biodomain knowledge graphs.

| biological domain | nodes (mRNA) | edges (mRNA) | nodes (proteins) | edges (proteins) |
| --- | --- | --- | --- | --- |
| APP Metabolism | 147 | 316 | 87 | 154 |
| Apoptosis | 958 | 10963 | 451 | 3261 |
| Autophagy | 505 | 2665 | 304 | 1096 |
| Cell Cycle | 728 | 6561 | 337 | 1779 |
| Endolysosome | 1096 | 8909 | 665 | 3874 |
| Immune Response | 1507 | 16354 | 688 | 4683 |
| Lipid Metabolism | 1671 | 13515 | 891 | 4366 |
| Metal Binding and Homeostasis | 801 | 4822 | 441 | 1398 |
| Mitochondrial Metabolism | 1273 | 7092 | 858 | 3493 |
| Myelination | 189 | 348 | 120 | 132 |
| Oxidative Stress | 346 | 2080 | 185 | 637 |
| Proteostasis | 2675 | 29812 | 1477 | 13323 |
| Structural Stabilization | 2245 | 23850 | 1297 | 12163 |
| Synapse | 2067 | 21056 | 1234 | 9686 |
| Tau Homeostasis | 45 | 90 | 41 | 81 |
| Vasculature | 756 | 7098 | 356 | 2008 |

### Modeling framework for multi-omics integration

In this work, we proposed an end-to-end framework for supervised integration of incomplete multi-omics data. Our modeling framework comprised two key components: 1) graph neural network-based feature extractors and 2) feature alignment as well as set transformer-based feature integration among modalities.

### Graph Neural Network-based feature extractors

Let 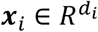 be the set of features for the *i*^*th*^ modality and 𝒢be an undirected graph with *d*_*i*_ nodes, with each node representing a feature in the *i*^*th*^ modality. 𝒢 may be constructed empirically via binarizing the matrix of correlation coefficients among features or may represent prior knowledge on the space of features defining known pairwise relationships. Let ℰ be the list of edges in 𝒢 and 𝒲 be an optional list of corresponding vector-valued edge weights such that |𝒲| *=* |ℰ|. Given a sample of ‘omics measurements ***x***_*i*_ and an associated graph topology 𝒢, we set up a graph neural network (GNN) and learned ***z***_*i*_ *= g*(***x***_*i*_, ℰ, 𝒲), where ***z***_*i*_ ∈ *R*^*m*^ is a vector of low-dimensional embeddings. A schematic of the GNN-based feature extractor is shown in Figure 7.

**Figure 7:**
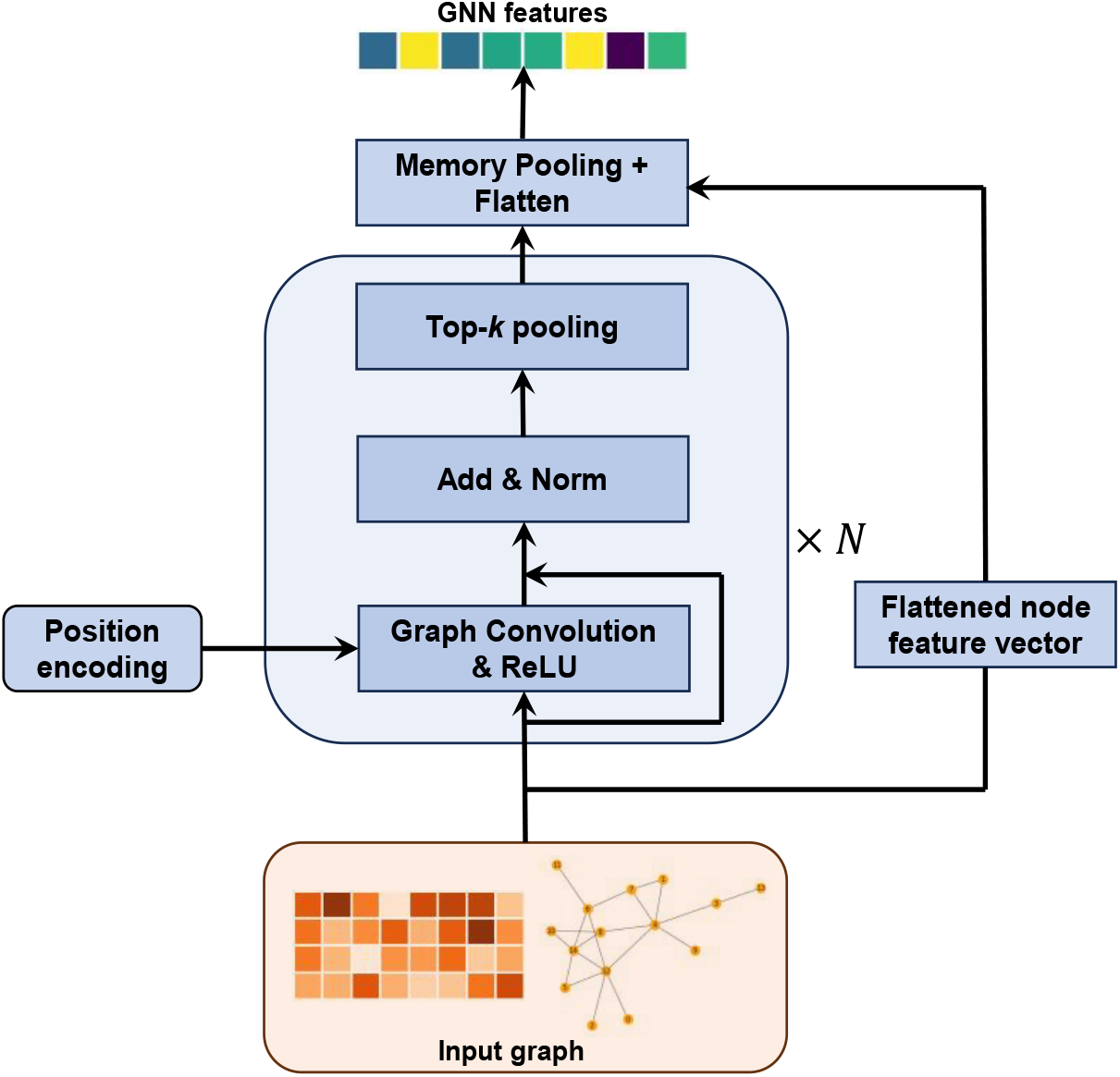
Schematic of GNN-based feature extractor. The feature extractor comprised of a stack of graph convolution blocks, a memory pooling layer and a residual connection between the original input node features and the output of the memory pooling layer.

First, we constructed a positional encoding of *d*_*i*_ features into *k* (*k <d*_*i*_) learnable communities using a standard lookup table embedding function in *Pytorch* which were softmax-normalized. The input node feature vector ***x***_*i*_ to *g* was linearly transformed by using the *k* communities as bases, followed by passing through a stack of graph convolution blocks. Each block (round-corner rectangular box in Figure 7) comprised three sub-blocks. The first sub-block was a message passing graph convolution layer, followed by a ReLU activation. The second sub-block was a residual connection adding input node features to the output of ReLU, followed by a batch normalization. The third sub-block was an optional top-*k* pooling layer [60], which we applied to mask non-informative nodes when input graphs were large. Specifically, the parameters in the top-*k* pooling layers were set so that the graphs before the memory pooling layer contained at most 300 non-masked nodes. Input node features which passed through *N* graph convolution blocks were transformed into latent node features 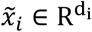. The associated graph topology remained unchanged since we did not employ edge-updating in our model. The transformed node feature vector 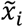 passed through a memory pooling layer [61] which learned a coarse graph representation through soft cluster assignments. Subsequently, 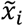 was reduced to 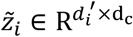, where *d*_*c*_ is the number of clusters and *d*_*i*_′ is the number of features in a cluster, which was then flattened. Lastly, a residual connection was employed adding the linearly transformed original input node feature vector to the flattened 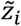, which was subsequently linearly mapped to *z*_*i*_ ∈ R^m^.

### Graph convolution layer

Let 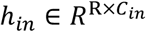 denote the input matrix to our graph convolution layer, where R was the number of nodes, and *C*_*in*_ represented the number of features in an input node. Let *C*_*out*_ be the number of features in an output node of a graph convolution layer. Suppose R nodes were mapped to k communities, and the softmax-normalized positional encoding were *p* ∈ R^R×k^. We first mapped the input node features linearly: *h*^′^ *= h*_*in*_W, where 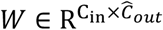 were learnable weights, 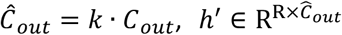. Linearly transformed node features *h*^′^ were then reshaped to 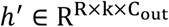. The positional encoding was then used to linearly weight node features along each output channel – 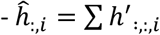 ⊙ unsquee*z*e(p), where ⊙ is the element-wise product operation and the summation was among k communities. The resulting node features were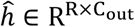. Finally, a message-passing update on the node features was performed:

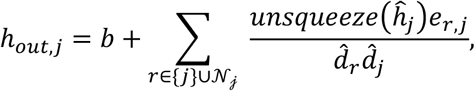

where 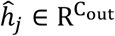 was the feature vector of the *j*^*th*^ node, 𝒩_𝒿_ was the set of indices of the neighbor nodes to node *j*, 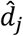 and 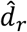 were the degrees of nodes *j* and *r* respectively, and 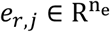 was a vector of *n*_*e*_ attributes for the edge connecting nodes *j* and *r*. Note that our graph convolution layer was an extension of the standard graph convolution of [62] where we allowed for the inclusion of positional embeddings and vector-valued edge attributes.

### Aligning embeddings among modalities

Before integration, we aligned embeddings among modalities by maximizing the similarities between modality embeddings. For samples with complete multi-omics data, we calculated the Pearson correlation between the two modality embeddings for each embedding dimension, and the similarity was calculated as the average of absolute Pearson correlations across embedding dimensions. For samples with incomplete multi-omics data (i.e., having only transcriptomic or proteomic measurements), we calculated the Pearson correlation for each pair of samples with the same class labels but different modalities, and the similarity was calculated as the average of absolute Pearson correlations across all possible sample pairs. The sum of the negative averages was included in the regularization loss terms.

### Integration using set transformers

Let *z*_*i*_ ∈ R^m^ be the GNN embeddings corresponding to the *i*^*th*^ ‘omics modality. We collected embeddings from GNNs corresponding to each individual modality into a set *Z =* (*z*_0_, *z*_1_, *⋯, z*_*k*_)^*T*^ ∈ R^(k*+*1)×m^, where *k*(≥ _1_) was the number of modalities and *z*_0_ ∈ *R*^*m*^ was a set of learnable parameters called the *class token*. We then used the standard transformer encoder architecture described in [63] to set up an integrative classifier. The prediction from the integrative classifier, *ŷ* ∈ R^C^ (where *C* was the number of classes) was given by -

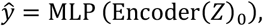

where Encoder(*Z*)_0_ ∈ *R*^*m*^ was the latent representation of the input learnable class token *z*_0_, and MLP(*⋅*): *R*^*m*^ *→ R*^*C*^ was a fully connected neural network which mapped the final class token representation to the target label space. The transformer encoder was a composition of *n* encoder blocks i.e., Encoder(*Z*) *= E*_*n*_ *∘ E*_*n−*1_*… ∘ E*_1_(*Z*). The sequence of operations within each encoder block was as follows:

1. Multi-head self-attention (MSA), residual connection and layer normalization (LN) [64] on the set of ‘omics tokens - *Z*^′^ *= LN*(*Z + MSA*(*Z*)),
2. Position-specific feedforward network followed by a residual connection and layer normalization (LN) - *Z*_*out*_ *= LN*(*Z*^′^ *+ FFN*(*Z*^′^)), where *FFN*(*⋅*):R^m^ *→*R^m^ was a one hidden layer MLP with ReLU activation and operated on each token individually, i.e., *FFN*(*Z*) *= FFN*(*z*_0_, *z*_1_, *…, z*_*K*_) *=* (*FFN*(*z*_0_), *FFN*(*z*_1_), *…, FFN*(*z*_*K*_))^*T*^).

### Full model architecture and training

Our full integrative multi-omics model was expressed as:

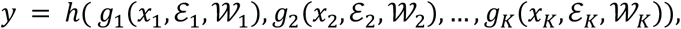

where, *h*(*⋅*) was a set transformer which integrated feature representations generated by the ‘omics GNNs *g*_*i*_*s*, ***x***_*i*_*s* were the node features for the *i*^*th*^ ‘omics modality and ℰ_*i*_, 𝒲_*i*_ were the edge index list and edge attribute list associated with the *i*^*th*^ modality. A schematic of the architecture is shown in Figure 1. Given multi-omics data for a given sample, measurements for each modality were processed through their respective GNN modules. The parameters of the GNN were trained by classifying the embeddings to the corresponding target labels using multi-layer perceptrons (MLP) or a set transformer which collected embeddings for all available modalities, integrated them, then made a prediction on the target label.

### Training with complete samples

We first describe the training of our model when the multi-omics data set had complete samples, i.e., we had measurements in all available ‘omics modalities for all patients. We split the total dataset into three folds, maintaining the ratio of sample labels in each fold as in the whole dataset. Samples from two of these splits were chosen as training samples while samples from the remaining split were used as validation samples. Since samples in our dataset had binarized labels (AD/control) we therefore used the binary cross entropy loss function. We set up our end-to-end integrative model as detailed in the previous sections and trained it using stochastic gradient descent optimization, specifically the Adam optimization method [65], using mini-batches of training data. The objective function we optimized was as follows –

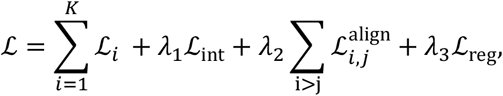

where *ℒ*_*i*_ was the loss incurred on the predictions made by the MLP classifier on the embeddings of the *i*^*th*^ modality, *ℒ*_int_ was the loss incurred on the predictions made by the integrative module, i.e., the set transformer, 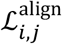 was the alignment loss between the *i*^*th*^ and *j*^*th*^ modalities, and *ℒ*_reg_ were norm penalties on the learnable parameters in the model. λ_1_, λ_2_,λ_3_ were trade-off hyper-parameters. Furthermore, we applied sample weights on our prediction losses (*ℒ*_*i*_ and *ℒ*_int_) to account for any potential class imbalance.

### Training with incomplete samples

To account for incomplete samples (i.e., samples with missing measurements in one or more modalities), we split our total training dataset into disjoint subsets based on modality representation. For our transcriptomics and proteomics datasets, we split the total dataset into three subsets – the set of common samples and two additional sets comprising samples having measurements in transcriptomics or proteomics alone. Each epoch of training for our integrative model now comprised a single epoch through each disjoint subset. The common sample subset was trained with the full loss function as described in the previous section. The loss function was modified for training data subsets with missing modalities. For a subset of the training data with measurements in a single modality only, the components *ℒ*_int_ and 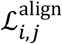 were set to 0. The order in which the disjoint data subsets were processed to update the model weights during training was randomized from epoch to epoch.

### Hyperparameter selection

As described in the previous section, we created 3 stratified splits of our dataset, from which we picked one fold for validation and used the remaining two for training. For any given combination of hyperparameters, we trained our models 3 times per split, each with a different random model weight initialization, and cross-validated on the validation dataset. In total, we trained the model 9 times with the same hyperparameters. We averaged the validation performance of our model over these 9 trials and reported it as the predictive performance corresponding to a given setting of hyperparameters. We performed a grid search over hyperparameters and picked the hyperparameters with the highest average validation accuracy over 9 trials.

For GNNRAI training, we conducted a non-exhaustive preliminary hyperparameter search, using 3 of the 16 biodomain datasets (autophagy, myelination and cell cycle), to establish baseline ranges for all hyperparameters and identify those to which model performance was most sensitive. During final model training we performed a smaller hyperparameter grid search over the set of critical hyperparameters identified during the preliminary stage. A summary of the hyperparameters explored during the final model training stage, along with a selection of the most critical fixed hyperparameters, are shown in Supplementary Table 1. Full details of the GNNRAI model structure were described in the section ‘Forward pass of the end-to-end GNNRAI model’ of supplementary material. For the benchmark MOGONET method, we adopted the same model architecture and training procedure described in [12]. A summary of the hyperparameter settings swept through for MOGONET training is shown in Supplementary Table 2.

### Integrated gradients

Let *x=* (*x*_1_, *x*_2_, *…, x*_*d*_)^T^ ∈ R^*d*^ be the input features to a deep learning model, *f*:R^*d*^ *→*R^*C*^, where *C* was the number of classes of the target label. Let *f*_*c*_(*x*) be the model output score for the _*c*_^*th*^ class. The components of the gradient vector, ∇_*x*_*f*_*c*_, represented the sensitivity of the class score to small perturbations of the input features, and the magnitude of its components may be interpreted as a proxy for the importance of input features. The integrated gradient attribution of the *i*^*th*^ input feature on the _*c*_^*th*^ class was defined as follows:

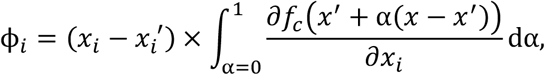

where *x*^′^ was a user-defined baseline input. The integrand represented the model evaluation at an input constructed by the linear interpolation between the baseline *x*^′^ and the true input *x*. The choice of the baseline was task dependent. For instance, in image classification tasks, where the input *x* is a tensor representing pixel intensities, it is customary to pick the zero tensor as the baseline, i.e., *x*^′^ *=* 0. In our multi-omics model, inputs were node features representing gene expression/protein abundance levels. We set our baseline as the average expression/abundance level in our training control samples. Features with large magnitude attribution scores on the disease class label (i.e., class label 1) were implicated as informative disease biomarkers.

### Integrated Hessians

Integrated Hessians for pairwise feature interactions [18] is a natural extension of the method of integrated gradients. Let ϕ_*i*_:R^*d*^ *→ R* be the integrated gradient attributions on the *i*^*th*^ input feature. Applying integrated gradient explanations on the function ϕ_*i*_(*x*) resulted in a new *d*-dimensional vector whose *j*^*th*^ component Γ_*i,j*_(*x*) *=* ϕ_*j*_(ϕ_*i*_(*x*)) explained the contribution of feature *x*_*j*_ to the model attributions on feature *x*_*i*_. We interpreted the quantity Γ_*i,j*_(*x*) as the interaction between features *i* and *j*. For *i* ≠ *j*, the feature interaction scores were given by:

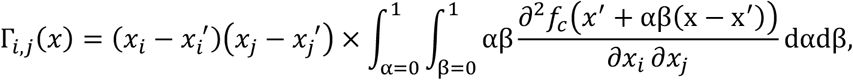

where *x*^′^ was a baseline input. The self-interaction term Γ_*i,i*_(*x*) was given by:

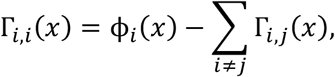

where the *i*^*th*^ feature integrated score ϕ_*i*_(*x*) represented the marginal contribution of *x*_*i*_ to model prediction. The self-interaction score Γ_*i,i*_(*x*) was, thus, defined as the difference between the marginal contribution of *x*_*i*_ and every pairwise interaction involving *x*_*i*_.

### Identify informative markers or marker interactions with specified false discovery rate (FDR)

To determine the importance score threshold above which a marker or a marker interaction was viewed informative, we adopted the permutation approach to compute the empirical FDR. Specifically, we randomly permuted the order of the ground truth labels to generate *B* permuted datasets of ground truth labels. We then trained the GNN model on each of the *B* permuted datasets and computed importance scores for markers using integrated gradients or marker interactions using integrated Hessians. Following the non-parametric procedure outlined in [66], we calculated the false discovery rate for a given threshold *d* as:

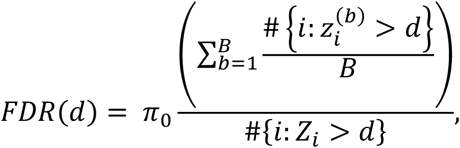

where, *Z*_*i*_ was the importance score of marker or interaction *i* in the unpermuted data, 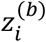 was the importance score of marker or interaction *i* in the *b*^*th*^ permutation, *π*_0_ was the prior probability that a marker or an interaction was uninformative. For AD study, *π*_0_ is close to 1, and we took the value of 0.97. The scores obtained under the null hypothesis (permuted data) could be used by different analyses from the same dataset – for instance, analyses from different random initializations and different training dataset folds. For an FDR threshold of ≤0.05, the estimated empirical FDR was confidently accurate for 100 to 200 permutations [24].

## Author Contributions

YL and GWC designed the project and analysis methods. HW, ZF, RT and YL developed the AI software, ZF processed data, ZF, RT and HW performed analyses using the AI framework, GAC, SK and GWC interpreted the analysis results in the context of AD. RT and YL drafted the manuscript, all authors revised the manuscript.

## Acknowledgments and Funding Sources

We would like to thank L Health for her advice on ROSMAP proteomics data quality control, and JC Wiley for his help in BD interaction interpretations. This study was supported by NIA grants R21 AG083299 and U54 AG065187. Data used in this project were obtained from the Accelerating Medicines Partnership Program for Alzheimer’s Disease (AMP-AD) Consortium members below:

## Religious Orders Study/Memory and Aging Project (ROSMAP)

We are grateful to the participants in the Religious Order Study and the Memory and Aging Project. This work was supported by the US National Institutes of Health (U01 AG046152, R01 AG043617, R01 AG042210, R01 AG036042, R01 AG036836, R01 AG032990, R01 AG18023, RC2 AG036547, P50 AG016574, U01 ES017155, KL2 RR024151, K25 AG041906-01, R01 AG30146, P30 AG10161, R01 AG17917, R01 AG15819, K08 AG034290, P30 AG10161, and R01 AG11101).

## Mount Sinai Brain Bank (MSBB)

This work was supported by grants R01AG046170, RF1AG054014, RF1AG057440, and R01AG057907 from the NIH/NIA. R01AG046170 is a component of the AMP-AD Target Discovery and Preclinical Validation Project. Brain tissue collection and characterization was supported by NIH HHSN271201300031C.

## Mayo RNAseq Study

Study data were provided by the following sources: The Mayo Clinic Alzheimer’s Disease Genetic Studies, led by Dr. Nilufer Ertekin-Taner and Dr. Steven G. Younkin, Mayo Clinic, Jacksonville, FL, using samples from the Mayo Clinic Study of Aging, the Mayo Clinic Alzheimer’s Disease Research Center, and the Mayo Clinic Brain Bank. Data collection was supported through funding by NIA grants P50 AG016574, R01 AG032990, U01 AG046139, R01 AG018023, U01 AG006576, U01 AG006786, R01 AG025711, R01 AG017216, R01 AG003949, NINDS grant R01 NS080820, CurePSP Foundation, and support from Mayo Foundation. Study data include samples collected through the Sun Health Research Institute Brain and Body Donation Program of Sun City, Arizona. The Brain and Body Donation Program is supported by the National Institute of Neurological Disorders and Stroke (U24 NS072026 National Brain and Tissue Resource for Parkinson’s Disease and Related Disorders), the NIA (P30 AG19610 Arizona Alzheimer’s Disease Core Center), the Arizona Department of Health Services (contract 211002, Arizona Alzheimer’s Research Center), the Arizona Biomedical Research Commission (contracts 4001, 0011, 05-901, and 1001 to the Arizona Parkinson’s Disease Consortium), and the Michael J. Fox Foundation for Parkinson’s Research.

## Data Availability

All the data analyzed in the article were downloaded from the AD Knowledge Portal https://adknowledgeportal.synapse.org [67]. The ROSMAP, MSBB, and Mayo bulk brain RANSeq data were from https://www.synapse.org/Synapse:syn30821562; the ROSMAP bulk brain TMT proteomics data were from https://www.synapse.org/Synapse:syn31534849; and the MSBB bulk brain TMT proteomics data were from https://www.synapse.org/Synapse:syn52331749. Alzheimer’s disease biodomain network files can be found at https://www.synapse.org/Synapse:syn51739831.

## Code Availability

Code will be made publicly available upon publication.

