## Supplementary File 1 for "An explainable graph neural network approach for effectively integrating multi-omics with prior knowledge to identify biomarkers from interacting biological domains"

### Supplementary material

#### Forward pass of the end-to-end GNNRAI model

We set up our end-to-end model with 2 graph convolutional blocks in the GNN feature extractor and 2 encoder layers in the set transformer integration module. For a single modality, the forward pass through the GNN was as follows:

Input graph (node feature matrix, edge index matrix, edge weight matrix).

1. Graph convolutional block 1 (GC1)
    i. Graph convolutional layer (as described in methods section)
    ii. ReLU activation
    iii. Residual connection from input to GC1
    iv. Batch normalization
    v. (Optional) TopK Pooling
2. Graph convolutional block 2 (GC2)
    i. Graph convolutional layer
    ii. ReLU activation
    iii. Residual connection from input to GC2
    iv. Batch normalization
    v. (Optional) TopK Pooling
3. Memory pooling
4. Residual connection from input node feature matrix to the output of step 3
5. Fully connected layer+ReLU+dropout+batch normalization
6. Fully connected prediction layer

The output of the forward pass through the GNN was to generate logit predictions on the basis of a modality which was used to compute the single modality component of the total loss as described in the Methods section. The result of step 5 in the above description was a low-dimensional feature representation for the given modality. All such feature representations were aggregated, from individual modalities, into a set of feature representations (or tokens) to be passed into the set transformer. The sequence of operations in the set transformer were as follows:

1. Encoder layer 1 (E1)
    i. Multi-head self-attention
    ii. Residual connection
    iii. Layer normalization
    iv. Position-wise feedforward layer
    v. Residual connection
    vi. Layer normalization
2. Encoder layer 2 (E2)
    i. Multi-head self-attention
    ii. Residual connection
    iii. Layer normalization
    iv. Position-wise feedforward layer

  v. Residual connection
  vi. Layer normalization
3. Fully connected layer on class token representation from previous step.
4. ReLU activation
5. Dropout
6. Fully connected prediction layer

**MOGONET model structure and training**
We followed the same architecture for MOGONET as described in [1] – three graph convolutional network (GCN) layers followed by a view correlation discovery network (VCDN). We also followed the same training procedure as specified in [1] – 'Omics specific GCN feature extractors are pretrained initially and a secondary end-to-end training stage is performed with the GCN and VCDN networks combined'.

**Supplementary Table 1:** Summary of scanned and critical fixed hyperparameters in GNNRAI training.

| **Hyperparameter** | **Values** |
|---|---|
| Graph embedding dimension | 16 |
| Batch size | 16 |
| Learning rate | $2.5\times10^{-3}$ |
| Number of clusters in memory pooling layer | 40, 80 |
| L1 regularization penalty in classifier | 0, $1\times10^{-3}$ |
| L2 regularization penalty in classifier | 0, $1\times10^{-3}$ |

**Supplementary Table 2**. Hyperparameters swept through for tuning the benchmark MOGONET model.

| **Hyperparameter** | **Values** |
|---|---|
| Number of edges per node, $k$ | 2, 3 |
| Number of hidden units in 1st two GCN layers, $h_1$ | 200, 400 |
| Number of hidden units in final GCN layer, $h_2$ | 100, 200 |
| Pretraining learning rate | $10^{-4}$, $10^{-5}$ |
| Learning rate for GCN layers during end-to-end training stage | $5\times10^{-4}$, $5\times10^{-5}$ |
| Learning rate for VCDN layer during end-to-end training stage | $1\times10^{-3}$, $1\times10^{-4}$ |
